# A machine learning based approach towards high-dimensional mediation analysis

**DOI:** 10.1101/2022.10.10.511329

**Authors:** Tanmay Nath, Brian Caffo, Tor Wager, Martin A. Lindquist

## Abstract

Mediation analysis is used to investigate the role of intermediate variables (mediators) that lie in the path between an exposure and an outcome variable. While significant research has focused on developing methods for assessing the influence of mediators on the exposure-outcome relationship, current approaches do not easily extend to settings where the mediator is high-dimensional. These situations are becoming increasingly common with the rapid increase of new applications measuring massive numbers of variables, including brain imaging, genomics, and metabolomics. In this work, we introduce a novel machine learning based method for identifying high dimensional mediators. The proposed algorithm iterates between using a machine learning model to map the high-dimensional mediators onto a lower-dimensional space, and using the predicted values as input in a standard three-variable mediation model. Hence, the machine learning model is trained to maximize the likelihood of the mediation model. Importantly, the proposed algorithm is agnostic to the machine learning model that is used, providing significant flexibility in the types of situations where it can be used. We illustrate the proposed methodology using data from two functional Magnetic Resonance Imaging (fMRI) studies. First, using data from a task-based fMRI study of thermal pain, we combine the proposed algorithm with a deep learning model to detect distributed, network-level brain patterns mediating the relationship between stimulus intensity (temperature) and reported pain at the single trial level. Second, using resting-state fMRI data from the Human Connectome Project, we combine the proposed algorithm with a connectome-based predictive modeling approach to determine brain functional connectivity measures that mediate the relationship between fluid intelligence and working memory accuracy. In both cases, our multivariate mediation model links exposure variables (thermal pain or fluid intelligence), high dimensional brain measures (single-trial brain activation maps or resting-state brain connectivity) and behavioral outcomes (pain report or working memory accuracy) into a single unified model. Using the proposed approach, we are able to identify brain-based measures that simultaneously encode the exposure variable and correlate with the behavioral outcome.

**HIGHLIGHTS:**

- Current methods for assessing mediation do not easily extend to high dimensions
- We introduce a new approach for performing high-dimensional mediation analysis
- Links high-dimensional mediator to path analysis model via machine learning algorithm
- Method illustrated using data from two fMRI studies

## II. INTRODUCTION

A frequent occurrence in biological, mechanical, and information systems alike is that the relationship between two variables *x* and *y* is transmitted through a third intervening variable, or *mediator, m*. An example of such a relationship is illustrated in the three-variable path diagram depicted in Figure 1A. For example, exposure to a drug may cause a clinical benefit via its effects on brain neurotransmitter levels. Solar energy may power an electric motor via an intermediate transformation to energy by a solar cell. Changing the position of an advertisement on a web page may influence sales of the advertised product via the position’s intermediate effects on people’s attention to the ad. In all these cases, estimating how much of the total effect of the exposure (or *initial variable, x*) on the outcome (or *dependent variable, y*) is transmitted through the mediator can help explain how the exposure influences the outcome, and thus under what conditions the relationship is likely to occur.

**FIG. 1.**
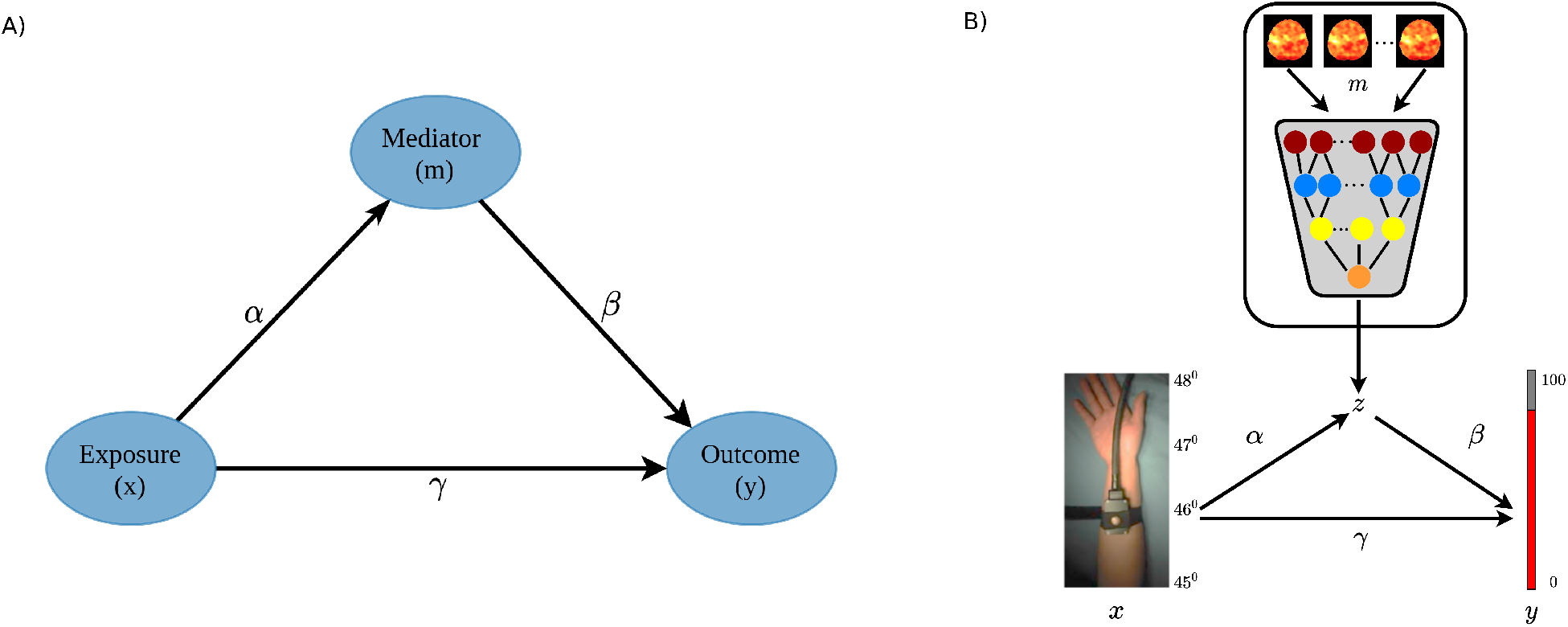
(A) An overview of the standard three-variable mediation model. The variables *x, m*, and *y* are all scalars along with the associated path coefficients, *α, β*, and *γ*. (B) Schematic representation of the proposed mediation analysis framework using deep learning. Here a deep learning model links the high-dimensional mediators (brain activation maps) to a standard path analysis model used to access mediation. The output of the deep learning model is a latent intermediate mediating variable between in the input stimulus intensity(*x*) and the reported pain (*y*). The goal is to evaluate whether there is a significant indirect effect *αβ*.

The concept of mediation has been a staple in the behavioral sciences [1] for a century, and a linear model version of mediation analysis was popularized in the psychometric and behavioral sciences literature several decades ago [2, 3]. This framework has since been widely used in the social and behavioral sciences [4], economics, decision and policy making [5, 6], epidemiology [7], neuroscience [8, 9], and beyond. It has also been extended to use estimates based on modern causal modeling frameworks [10–14].

In most applications, the mediator variable is either univariate [15, 16] or low-dimensional, meaning that there are typically only one or a few mediating variables in the model [17–19]. In practice, in many psychological, behavioral, and biological systems, there are many potential mediators, and these can be highly correlated. For example, the effects of surgery on post-operative pain may be mediated by a complex pattern of correlated gene expression changes in immune cells. The effects of an advertisement campaign on sales may be mediated by a complex pattern of measurable user data. Similarly, the effects of a hot stimulus on reported pain might be mediated by a complex pattern across inter-correlated brain regions. When the mediator space is high-dimensional, with larger numbers of mediators and multi-colinearity among them, estimating individual path coefficients in the standard way is not feasible. Potential reasons include difficulties modeling the appropriate relationship between variables in this setting [20], and the fact that standard estimation procedures become unstable when the number of mediators is much larger than the number of observations. However, in many cases, including those above, it may be useful or even preferred to assess the effects of a pattern across mediating variables of the same type in aggregate, without attempting to disentangle the unique causal effects of any single one. For example, the unique effect of each of 10,000 gene expression measures on post-operative pain may be difficult or impossible to estimate adequately, but a pattern that constitutes some function across the set of inter-correlated variables (e.g., a weighted average) may be both possible to estimate precisely and useful for both predictive and explanatory purposes. Such summaries are increasingly popular in genetics, neuroimaging, -omics, and beyond [21–23]. In genetics, for example, it is now possible to measure ∼ 1 million inter-correlated single-nucleotide polymorphisms, which individually explain *<* 1 percent of the variance in phenotypes at best, but in aggregate can often explain much more variance. These pattern-based models have enjoyed wide applicability in machine learning, but have seldom been extended to mediation tests. Thus, with the recent growth in the number of new applications collecting data on massive numbers of variables (e.g., brain imaging, genetics, epidemiology, and public health studies), it has become important to develop mediation analysis in high-dimensional settings.

As a motivating example that we continue throughout the remainder of this paper, consider the study of human brain function using functional magnetic resonance imaging (fMRI) data. Here researchers are interested in understanding the role of distributed brain measures acting as potential mediators on the relationship between an exposure (or treatment) variable and certain cognitive (or outcome) variables [19, 24–30]. In this context, the mediator can be a high-dimensional image (e.g., a 3-dimensional structural brain image or brain activation map) or a set of measures of functional connectivity (e.g., a 2-dimensional connectivity matrix), while both the exposure and outcome variables are univariate. For instance, [31] uses functional connectivity to perform mediation analysis and suggest that prenatal exposure to crime is associated with weaker neonatal limbic and frontal functional brain connectivity.

Standard mediation techniques will not be directly applicable in these settings, and new approaches are required. [32] proposed an early approach based on expressing the multivariate images using summary measures upon which standard mediation analysis was performed. Another early approach, “mediation effect parametric mapping” [26– 28], sought to investigate univariate mediators at each spatial location (voxel). However, this ignores the inherent relationship between voxels, instead identifying a series of univariate mediators. More recently, a number of approaches have sought to explicitly derive optimized, multivariate linear combinations of the high-dimensional mediators. [33] proposed a transformation model using spectral decomposition where mediation effects were estimated by placing the univariate transformed mediators into a series of regression models. A related approach, denoted the “principal directions of mediation” (PDM) [34, 35], decomposed high dimensional mediators into multiple orthogonal mediators that together mediate the effect of an exposure variable on the outcome. The method was applied to fMRI data and used to identify brain regions that mediate the relationship between a thermal stimulus and reported pain [35]. Finally, [36] proposed a sparse principal components approach towards high-dimensional mediation analysis.

In this paper we introduce a novel machine learning based method for identifying high dimensional mediators. Our proposed approach links the high dimensional mediators (e.g., brain activation maps or resting-state functional connectivity) to a standard path analysis model through a machine learning model (e.g., deep learning or support vector regression); see Figure 1B. Our proposed algorithm iterates between using a machine learning model to map the high-dimensional mediators *m* onto low dimensional mediators *z*, and using the predicted values as input in a standard three-variable mediation model. Importantly, the true value of *z* is latent, and the machine learning algorithm is trained to maximize the likelihood of the underlying mediation model, rather than based on directly predicting *z*. Our proposed approach uses an iterated maximization algorithm that alternates between fitting the machine learning algorithm and the mediation model. Thus, the approach provides a means of linking exposure variables, high-dimensional brain measures, and behavioral outcomes into a single unified model. Importantly, our proposed algorithm is flexible enough to allow researchers to ‘plug in’ various different types of machine learning algorithms, depending on the type of data assumed to mediate the relationship between exposure and outcome. In this work we explore a variety of such plug-ins, including a deep learning model, a shallow learning model, support vector regression, and a connectome-based predictive model [37]. Research on high-dimensional mediation analysis is in its infancy and this is to the best of our knowledge the first application of deep learning to the field.

We illustrate the performance of the proposed method through a simulation study and application to two different fMRI datasets. In the first application, we use data from eight different heat pain studies (N=284) to investigate the role of brain mediators on the generation of pain experience. Here a series of thermal stimuli were applied at various temperatures to each subject. In response, subjects gave subjective pain ratings at a specific time point following the offset of the stimulus. During the course of the experiment, brain activity in response to the thermal stimuli was measured across the entire brain using fMRI. The goal is to determine brain regions whose activity level act as potential mediators of the relationship between temperature and pain rating. In this application we use the proposed algorithm together with a deep learning model. Seven out of the eight studies (N=209) were used as training data, and the final study (N=75) was used as test data. Here the model parameters estimated in the training data are used to validate model performance in new set of individuals. Figure 2A provides an overview of the proposed setup. Importantly, the test data set not only included heat pain stimuli, but also physically and emotionally aversive sounds, providing a test of whether brain mediators of pain are specific to pain or general across pain and aversive sounds. While the derived mediators should generalize to different pain data sets, they are not expected to mediate the relationship between sound levels and perceived sound intensity. We benchmark the performance of our approach against [35], which used the same data to find high-dimensional brain patterns that mediate pain using the linear PDM approach, and mass-univariate mediation effect parametric mapping.

**FIG. 2.**
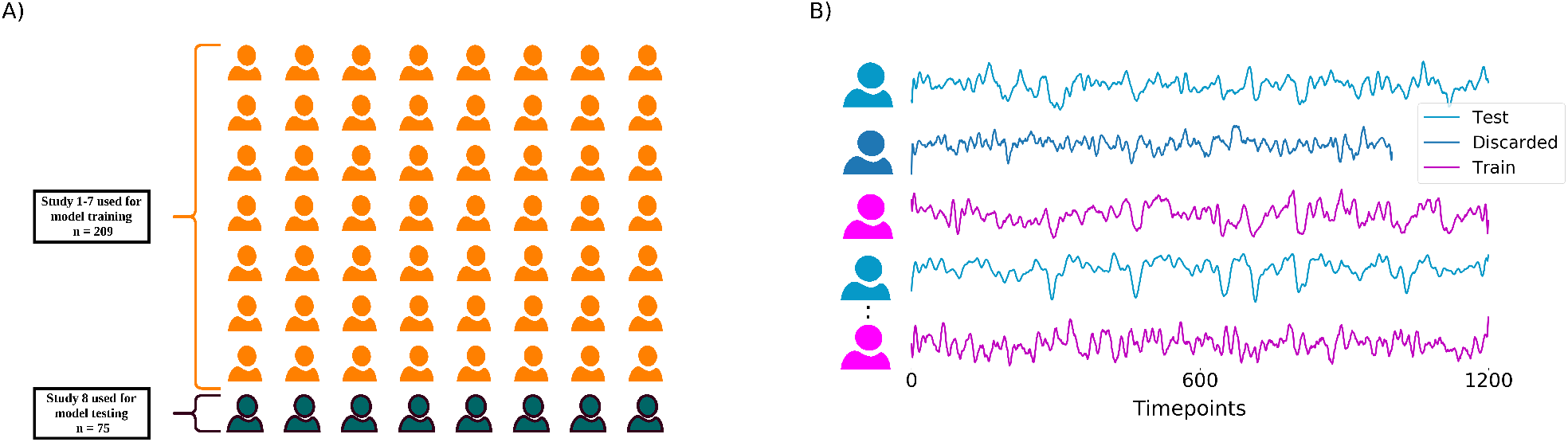
(A) Overview of Pain data. Studies 1 − 7, which comprise of 209 subjects and 13,372 trials, are used to train the deep learning model. The model is tested independently on Study 8 which consists of 75 subjects and 2,296 trials. Additionally, Study 8 not only includes the type of thermal pain stimuli used to train the model, but also aversive sounds. We tested our model and hypothesized that the estimated brain mediators of pain should generalize to the new pain dataset, but not to the sound dataset. (B) Overview of the HCP dataset. We used rs-fMRI data with *LR* polarity (*rf MRI*_*REST* 1_*LR*) from the HCP 900 release to investigate the relationship between working memory accuracy, measured using performance on an N-back task, and fluid intelligence. We excluded subjects with missing time points. We used 70% of the selected subjects for training and 30% for testing the model

In the second application, we use behavioral and resting-state fMRI (rs-fMRI) data from the Human Connectome Project (HCP) 900 release [38] to investigate the relationship between fluid intelligence and working memory, measured using performance on an N-back task. In particular, we sought to explore whether resting-state brain connectivity measures mediated the relationship between these two variables. In this application we use the proposed algorithm together with a connectome-based predictive model [39]. For each subject we extracted the mean time series from 268 regions of the Shen atlas [40], and computed a connectivity matrix where each element represents the Pearson correlation between the time series from two regions. We sought to investigate whether the elements of the correlation matrix mediated the relationship between intelligence and accuracy. In total we had 798 subjects with complete data, where 70% were used for training the model and 30% for testing. Figure 2B provides an overview of the proposed setup. These two examples illustrate the ability of our approach to handle different types of data and utilize different types of models, highlighting the strength and flexibility of the proposed approach.

## III. METHODS

### A. Mediation model

Mediation analysis is an analytic technique used to make statistical inferences on the path coefficients (see Figure 1), particularly on the proportion of the total effect of *x* on *y* is mediated through *m*. The effects of the exposure on the outcome are decomposed into separable direct and indirect effects, representing the influence of the variables *x* on *y* unmediated and mediated by *m*, respectively. Using the notation in Figure 1, the indirect effect is given by the product of the coefficients *α* and *β*, and the direct effect by the coefficient *γ*. Together, their sum represents the total effect of *x* on *y*.

Here we introduce our machine learning-based method for identifying high-dimensional mediators; see Figure 1B. For *i* = 1, …, *n*, where *n* denotes the number of trials, let *x*_*i*_ and *y*_*i*_ denote the univariate exposure and outcome variables, respectively, and let *m*_*i*_ be a high-dimensional object consisting of *p* elements where *p >> n*. Further, let Φ(.) denote an arbitrary machine learning model that operates on the variables *m*_*i*_. Our proposed approach takes the output of the algorithm *z*_*i*_ = Φ(*m*_*i*_) and places it into a standard 3-variable mediation model together with *x*_*i*_ and *y*_*i*_. Importantly, we consider the true value of *z*_*i*_ to be a latent variable, and the machine learning model is instead trained to maximize the likelihood of the underlying mediation path analysis model (see (3)), rather than based on predicting *z*. Our proposed approach achieves this goal by using an iterated maximization algorithm that alternates between fitting the machine learning algorithm and the mediation model. Thus, all three variables *x*_*i*_, *m*_*i*_, and *y*_*i*_ are part of the loss function. To elaborate, we assume that the relationship between the variables is given by two sets of equations. First, the mediator model links the exposure to the output of the machine learning model as follows:

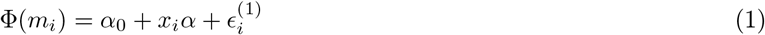

where *α*_0_ is the intercept, *α* is the coefficient describing the exposure-to-mediator relationship, and the error term 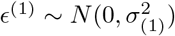. Second, the outcome model links the exposure and the output of the machine learning model to the outcome as follows:

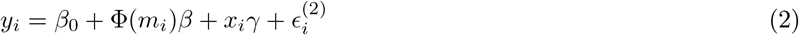

where *β*_0_ is the intercept, *β* is the coefficient representing the mediator-to-output relationship, *γ* is the direct effect of the exposure on the outcome, and the error term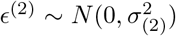. Once the parameters have been estimated we can express the total effect *τ* as the sum of the direct and indirect effects as follows: *τ* = *γ* + *αβ*. This is equivalent to the decomposition obtained in a standard univariate mediation analysis [2], and one can investigate whether there exists a significant mediation effect by testing: *H*_0_ : *αβ* = 0.

We propose to jointly fit all model parameters, including those in the machine learning model, through a single unified modeling approach. Combining the error terms from equations (1) and (2), the global loss function contribution over all observations is given by:

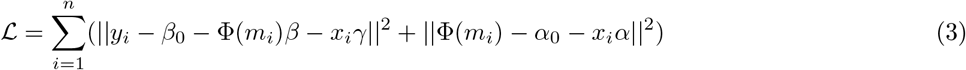

The solution to the global loss function corresponds to the maximum likelihood estimate of the three variable path model under normality assumptions.

In order to estimate the model parameters we propose an iterative algorithm that alternates between fitting the machine learning model Φ and the three-variable mediation model. Let us begin by assuming that the parameters *α*_0_, *β*_0_, *α, β*, and *γ* are known, the goal is to find an optimal solution for equation (3). Let *e*_*i*_ = *y*_*i*_ − *β*_0_ − *x*_*i*_*γ* and *h*_*i*_ = *α*_0_ + *x*_*i*_*α*. Then, keeping track of only those terms that involve Φ and completing the square, equation (3) becomes:

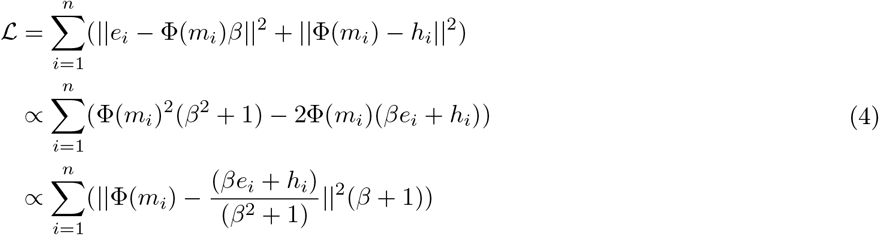

Under the assumption that *β, e*_*i*_ and *h*_*i*_ are known, minimizing the loss L is now equivalent to minimizing

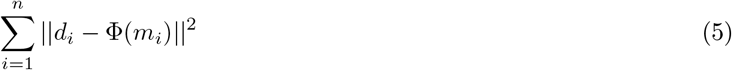

where 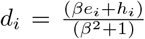This provides an appropriate loss function to fit the machine learning algorithm. Under the assumptions discussed above, the values of *d*_*i*_ are known and thus the model can be fit using standard estimation techniques.

Next, under the assumption that *z*_*i*_ = Φ(*m*_*i*_) is known, the parameters *α*_0_, *β*_0_, *α, β*, and *γ* can be estimated using a standard 3-variable path analysis model [2]. This involves fitting the regression models:

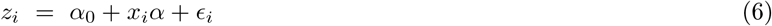

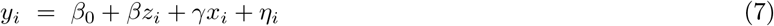

Solving the equations provides estimates of both the direct effect *γ* and the indirect effect *αβ* used to assess mediation. Initial estimates of *z*_*i*_ are computed using a given starting value for the the machine learning algorithm Φ. In the proposed framework the sign of *z*_*i*_ is not identifiable. Hence, we fix the sign so that the correlation between *z*_*i*_ and *y*_*i*_ is positive across observations *i* to simplify interpretation. A similar constraint is used when estimating the principal directions of mediation [35] and in independent components analysis (ICA). In addition, we normalize the variable *z*_*i*_ to avoid overshooting or shrinking of the prediction while iteratively minimizing the loss function expressed in Eq. 5. Thus, *z*_*i*_ is known up to sign and scale. Importantly, neither of these constraints affect the total amount of variance explained by the mediators. Using the exposure variable *x*_*i*_, outcome *y*_*i*_ and mediator *z*_*i*_, we fit the standard path analysis model to obtain the coefficients *α*_0_, *β*_0_, *α, β, γ*. Thereafter, we update the parameters of the machine learning model by fitting the model using *d*_*i*_ as the outcomes and *m*_*i*_ as the predictors. The proposed approach utilizes an iterative maximization algorithm that alternates between fitting the machine learning algorithm and the path analysis model. The pseudocode for the algorithm is described in Algorithm 1.

It is important to note that Algorithm 1 is agnostic to the choice of machine learning model. In this work, using simulated data, we show the flexibility of Algorithm 1 using: (1) a deep learning model; (2) a shallow learning model; and (3) support vector regression. As a demonstration, we apply the same deep learning model to the pain data to determine brain regions mediating the relationship between input stimuli and pain ratings. Additionally, we apply a ridge regression connectome-based predictive model [39] to the HCP data to determine the functional networks that mediate the relationship between fluid intelligence and working memory accuracy. Below we describe each model in turn.

#### Algorithm 1

Block maximization algorithm

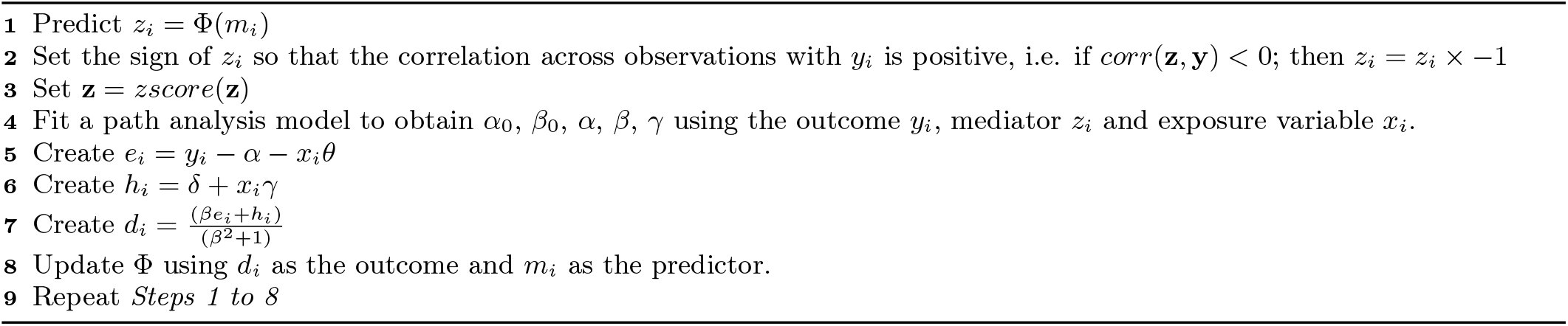

#### Deep learning model

We built a 3-dimensional convolutional neural network (CNN) based deep learning model that uses a residual architecture (ResNet) [41]. For the application to the pain data set, the input of the CNN consists of 3-dimensional volumes of size 91 × 109 × 91. Each volume corresponds to the brain activation map from a single trial. The CNN architecture consists of 5 residual blocks, each followed by a max pooling layer and a fully connected layer. The max pooling layer uses a stride of 2 with a kernel size of 3. Our model is inspired by a 3D-CNN based deep learning model used for brain age prediction [42]. However, our proposed model differs in two key areas. First, since we want a generalized model, we did not include information about sex and scanner type to the final layer. Second, we replaced the Batch re-normalization layer with a Batch normalization layer. The convolutional part of the CNN reduces the input image from dimensions 91 × 109 × 91 to 128 feature maps of size 3 × 4 × 3. The model was trained by minimizing the mean absolute error (MAE) using Adam optimization. The final fully connected part uses these feature maps to predict the lower-dimensional mediators. A flowchart of the model is shown in Figure S1. The CNN architecture is implemented using Keras version 2.4.0 [43] and Tensorflow version 2.3.1 [44] as the backend. We fit the deep learning models on the Oracle cluster using NVIDIA V100 Tensor core GPU. For the simulation study, the model was altered based on the dimensions of the input images; see below for a thorough description of the simulations performed.

#### Shallow learning model

We used a shallow CNN-based learning model with fewer layers than the deep learning model. The model consists of two convolutional layers, each with filter size 32 and 64 respectively, with kernel size 3 and using the rectified linear unit (ReLu) activation function [45]. The convolutional layers are followed by a max pooling layer with stride 2 and a dropout layer to reduce overfitting. Thereafter, a dense layer with filter size 128 and ReLu activation function is added followed by a dropout layer with keep rate equal to 0.5 and the final output layer with no activation function. Thus, the final layer performs a linear regression on the features of the hidden layers. Similar to the deep learning model, the MAE is used as the loss function and Adam optimization is used to ensure that the architecture converges. Further details about the training process is described in Section III D. The shallow learning model is used only in the simulation study for comparison purposes and to demonstrate the flexibility of our proposed approach.

#### Support vector regression

We used a non-linear support vector regression (SVR) using a radial basis function kernel. The python library scikit-learn [46] was used to implement the SVR and its regularization parameter was set to 1. Similar to the shallow learning model, SVR is only used in the simulation study for comparison purposes and to demonstrate the flexibility of our proposed approach.

#### Ridge regression connectome-based predictive modeling (rCPM)

We used a ridge regression connectome-based predictive model [39], which is an approach that has proven useful for developing predictive models of brain-behavior relationships from connectivity data. Here the features are obtained from a connectivity matrix where the edge of the matrix represents the Pearson correlation between the time series from two regions. Each edge in the connectivity matrix is related to the behavioral measures using a form of linear regression and a set of edges are selected using a significance test. Thereafter, a multivariate ridge regression model is fit to evaluate the brain-behavior relationship using the selected edges. The hyper-parameter corresponding to regularization strength is tuned using a 5-fold cross-validation grid search strategy which allows for an exhaustive search over the specified grid of parameters values (*λ* is allowed to take 100 evenly spaced values between 5*e* 3 to 5*e*9).

### B. Simulations

We performed three simulations to evaluate the performance of the proposed algorithm. We simulated a situation in which the latent mediator scores *z* are a complex, nonlinear function of an observed set of mediator variables. To accomplish this, in our simulations, we embedded the mediator in an image whose pixels represent the values of handwritten digits. In order to create a simulated dataset, we first fixed values of *α*_0_, *α, β*_0_, *β*, and *γ* for each simulation as described below. Next, we randomly generated input data *x* using a standard normal distribution with mean 0 and standard deviation 1. Thereafter, we used the input data as an explanatory variable in the linear regression model:

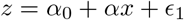

where *ϵ*_1_ ∼ *N* (0, 1). This allowed us to generate the low-dimensional mediators *z*. In order to create a high-dimensional set of mediators *m* that encode this information in a nonlinear fashion, we computed the cumulative distribution function of *z*, which gives us a value between 0 and 1. Next, we took the first 4 digits after the decimal point and found images of these digits in the MNIST dataset [47]. We concatenate the 4 images into a larger image to create the high dimensional mediators. Next, we simulated the outcome using the linear regression model:

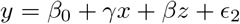

where *ϵ*_2_ ∼ *N* (0, 1). Steps for creating the dataset are summarized in Figure S1.

Using this framework for data generation, we performed three simulations, where for each, we evaluated the performance of Algorithm 1 using three different machine learning models: (1) a deep learning model; (2) a shallow learning model; and (3) support vector regression. The details of the implementation of each machine learning model are described above. The input to each of the models are the high-dimensional mediators *m* (computed using simulated data as illustrated in Figure S1) and *d*. The output is the model with tuned hyper-parameters that will be used for estimating the parameters of the path analysis model expressed in Algorithm 1.

In Simulation 1, we sought to evaluate how the sample size effects the ability to estimate the parameters of the mediation model shown in Figure S1. We varied the number of observations (subjects) while keeping the dimensions of the mediator constant. We used the MNIST data with image size 28 × 28 pixels, thereby creating a high-dimensional mediator with dimensions 28 × 112, for 100, 500 and 1000 observations. The model parameters were set to *α* = 0.2, *α*_0_ = 0.1, *β*_0_ = 6, *β* = 4, *γ* = 5, and *αβ* = 0.8.

In Simulation 2, we sought to evaluate how the size of the high-dimensional mediator impacts the ability to estimate the parameters of the mediation model. We fixed the number of observations to 1000, but varied the dimensions of the MNIST data. We scaled the MNIST images to 8 × 8, 32 × 32, 64 × 64 pixels, thus changing the dimension of the high-dimensional mediator variable to 8 × 32, 32 × 128, and 64 × 2 × 6. The values of the parameters *α, β, γ*, and *α×β* remained the same as in Simulation 1.

Finally, in Simulation 3, we sought to evaluate the performance of the model in a null-setting, where there is no significant mediation effect. We removed the link between the exposure and the mediator variable by setting the value of *α* to 0. The values of all other parameters remained the same as in Simulation 1. Similar to Simulation 1, we varied the number of observations (100, 500 and 1000 subjects) while keeping the dimensions of the simulated mediators constant (28 × 112).

For each simulation we fit the model for each of the three machine learning methods for 20 iterations. These iterations are used for estimating the coefficients *α, β, γ*, and the indirect effect *αβ*. It was noticed that the value of the coefficients converge in less than 5 iterations. This procedure was repeated 100 times for each model and simulation.

For comparison purposes, we also fit the PDM approach [34, 35] to the simulated data. This approach linearly combines information across images into a smaller number of orthogonal components that are chosen based on the proportion of the indirect effect that they explain. To facilitate comparisons with the proposed approach, we only use the first PDM which corresponds to the direction (or linear combination of features) that maximizes the proportion of the indirect effect explained. Subsequent PDMs, which maximize the remaining indirect effect conditional on being orthogonal to previous PDMs, are not used in this analysis.

### C. Experimental data

#### Participants

##### Pain Data

The data consisted of 284 healthy participants from eight independent studies [48–52] of thermal pain. The sample size in each study varied between *N* = 17 to *N* = 75 subjects. All participants were recruited from the New York City and Denver/Boulder areas and provided written informed consent. The institutional review board of Columbia University and the University of Colorado Boulder approved all studies. An online questionnaire, a pain safety screening form, and an fMRI safety screening form were used to determine the eligibility of all the participants. Any participant with psychiatric, physiological or pain disorders, neurological conditions, and MRI contraindications were excluded prior to enrollment. Additionally, participants were required to have at least 30 trials [35] with low variance inflation factors (< 3.5), non-missing ratings, and stimulation intensity data. Based on these criteria, 18 participants were excluded from Study 8.

##### HCP Data

The data consisted of subjects from the Human Connectome Project (HCP) 900 release [38] from the Washington University - University of Minnesota (WU-Minn HCP) Consortium. All participants gave full consent to the WU-Minn HCP Consortium, and research procedures and ethical guidelines were followed in accordance with Washington University institutional review board approval. Here resting state fMRI (rsfmri) data with *LR* polarity (*rf MRI*_*REST*1_*LR*) with 1200 time-points were used. Any subject with less than 1200 time-points or with missing data (i.e., with ‘nan’ values in the time series) were excluded from the analysis. Further, any participant with a missing fluid intelligence score or accuracy measure on the working memory task were also excluded. After excluding all such participants (n = 102 exclusions), 798 subjects remained and were included in our analysis.

#### Procedure

##### Pain Data

All participants received varying levels of thermal stimuli and rated their experienced pain while they underwent fMRI scanning. The number of trials, stimulation sites, rating scales, stimulus duration and intensities, inter-trial intervals varied across the studies, but were comparable; see [53] for further information. During fMRI scanning, the temperature of the heat stimulus (exposure variable) and pain rating (outcome variable) were recorded for each participant. Single trial brain activation maps were estimated using a general linear model (GLM) approach. In addition to the heat stimulus, participants in Study 8 also received an aversive sound stimuli during the fMRI scanning. The aversive sounds are taken from the the International Affective Digital Sounds database [54]. Example sounds include those of a knife scraping a plate (the single most aversive sound in the database) and emotionally aversive sounds like attacks, screaming and crying. Trials specific to aversive sounds were used to test the specificity of brain mediator patterns to thermal stimulus intensity and pain.

##### HCP Data

In addition to extensive MRI scanning, all HCP subjects performed a battery of cognitive tasks. Here we focus on measures of fluid intelligence and working memory accuracy. Fluid intelligence, a measure of higher order relational reasoning, was assessed using a 24-item version of the Penn Progressive Matrices test [55]. Working memory accuracy was measured using the mean accuracy across all conditions in an n-back task, described in detail in [56], and consisted of values between 0-100. During fMRI scanning, four 15-minute fMRI scans (runs) with a temporal resolution of 0.72 seconds and a spatial resolution of 2-mm isotropic were collected. Data from a single scan was used to create a resting-state connectivity matrix, described in more detail below.

#### fMRI data processing

##### Pain Data

Structural T1-weighted images were co-registered to the mean of the functional image. Thereafter, the registered image was normalized to MNI space using SPM(http://www.fil.ion.ucl.ac.uk/spm/). Studies 1 and 6 used SPM5, while SPM8 was used for all other studies. Following initial normalization, an additional normalization step based on the genetic algorithm-based normalization [57, 58] was performed in Studies 1 and 6. The first few volumes (ranging from 3-5) of each functional dataset was removed from the analysis to allow for image stabilization; see [53] for more detail. Mean and standard deviation of intensity values across each slice was used to identify outlier slices. Additionally, the Mahalanobis distance was computed for slice-wise mean and standard deviation of functional volumes. After false detection rate (FDR) correction for multiple comparisons, values with a significant *χ*^2^ value were considered as outliers. In total less than 1% of the total images were considered as outliers. The output of this procedure was included as nuisance covariates in subject-level models. Next, except for Study 8 (multiband data with a short TR of 480 ms), slice timing correction and motion correction was performed on the functional images using SPM. Functional images were warped to SPM’s normative atlas, interpolated to to 2 × 2 × 2*mm*^3^ voxels, and smoothed with an 8 mm FWHM Gaussian kernel.

For all studies except Studies 3 and 6, a single trial design and analysis approach was used to model the data by constructing a GLM design matrix with separate regressors for each trial [59, 60]. To model the cue and rating periods for each study, boxcar regressors were convolved with the canonical hemodynamic response function (HRF). Regressors for each trial, as well as several types of nuisance covariates were also included. Trial-by trial variance inflation factors (VIF) were calculated, and any trials with VIFs exceeding 2.5 were excluded from the analyses (VIF threshold for Study 8 was 3.5 as in the primary publication). For Study 1, global outliers (trials that exceeded three standard deviations above the mean) were also excluded, and a principal component based denoising step was employed during preprocessing to minimize artifacts. This generated single trial estimates that reflect the amplitude of the fitted HRF on each trial and represent the magnitude pain-period activity for each trial in each voxel. For Studies 3 and 6, rather than using a canonical HRF, single trial analyses were based on fitting a set of three basis functions. This allowed the shape of the modeled HRF to vary across trials and voxels. This procedure differed from that used in other studies because it maintains consistency with the procedures used in the original publications [58]. The pain period basis set consisted of three curves shifted in time and was customized for thermal pain responses based on previous studies [58, 61]. For Study 6, the pain anticipation period was modeled using a boxcar epoch convolved with a canonical HRF to estimate the cue-evoked responses. This epoch was truncated at 8 s to ensure that fitted anticipatory responses were not affected by noxious stimulus-evoked activity. Similar to other Studies, the nuisance covariates were included and trials with VIFs larger than 2.5 were excluded. In Study 6 trials that were global outliers (more than 3 standard deviations above the mean) were also excluded. The fitted basis functions from the flexible single trial approach were used to reconstruct the HRF and compute the area under the curve (AUC) for each trial and in each voxel. These trial-by-trial AUC values were used as estimates of trial-level pain-period activity. Together, these single trial maps of pain-period activity were used for model development and validation. The brain activation map for each participant was z-scored for each study. The final dimensions of the maps were 91 × 109 × 91. These maps were used as the high-dimensional mediators in our analysis.

##### HCP Data

For each subject, four 15 minute rs-fMRI scans with a temporal resolution of 0.72 seconds and a spatial resolution of 2-mm isotropic were available. We used the preprocessed and artifact-removed rs-fMRI data provided through the HCP900-PTN data release. This data has been extensively described in multiple other publications, so we only briefly discuss it here. The preprocessing pipeline followed the procedure outlined in [62]. Spatial preprocessing was applied using the procedure described by [63]. Independent component analysis (ICA), followed by FMRIBs ICA-based X-noisefier (FIX) from the FMRIB Software Library (FSL) [64], was used for structured artifact removal, removing more than 99 percent of the artifactual ICA components in the dataset.

Functional parcellation of each subject’s data was performed using the Shen atlas [40], which consists of 268 regions. For each region, the mean time series was extracted and shifted to 0 mean and unit variance. Any subject with less than 1200 time-points or with missing data (i.e., with ‘nan’ values in the time series) were excluded from the analysis. The Pearson correlation between each regions time course was computed, resulting in a 268 × 268 correlation matrix depicting functional connectivity between regions. Since these correlation matrices are symmetric, we vectorized the lower triangle of the matrix and used these values as the high-dimensional mediator in our analysis.

### D. Model fit and training procedure

The same general training procedure was used for both the simulated data and fMRI data, with the main difference lying in the number of iterations that were performed. For the simulated data Steps 1-8 of Algorithm 1 was iterated 20 times, while for the fMRI data it was only iterated 10 times to reduce computational burden.

In the simulation study, we evaluated the deep learning model, the shallow learning model, and support vector regression within our framework. Since the simulated data was created using MNIST data, all the layers of the deep and shallow learning models were constructed for 2D input data. For each simulation, 30% of the data was used as a validation data set, allowing us to judge how well the model generalized. The parameters of the mediation model *α, β* and *γ* were computed and compared with the ground truth value after the 20^*th*^ iteration. Both the deep and shallow learning models use the MAE as the loss function and the Adam optimization [65] method to ensure the architecture converges. The Adam parameters are set as follows: learning rate = 0.001, decay = 10^*−*6^, *β*_1_ = 0.9, *β*_2_ = 0.999, and batch size = 32. The model weights were initialized using the He initialization strategy [66] and a regularization parameter [67] *λ* = 5 ×10^*−*5^ is added to each trainable node in the CNN.

For the pain data, we combined our proposed algorithm with a deep learning model. In contrast to the stimulation study, the layers of the deep learning model were constructed for 3D input data. The first seven studies (N = 209) were used as training data, while the eighth (N=75) was used as the testing data [48–52]. Further, 30% of the training data were used as a validation set. The validation set is used to provide insight into whether or not the model is overfitting. To further check for overfitting, we stop training the model [68] if the validation error does not improve in 25 epochs and the weights with the lowest validation error were used for making the prediction in the test data. To evaluate the stability of the findings, we performed a leave-one-study out cross-validation where we alternated which of the eight studies were used as the validation dataset, while training on the remaining seven studies. In addition, we also ran multiple iterations of *k*-fold cross validation.

The input data consisted of a set of processed fMRI activation maps in response to the painful stimuli registered to MNI space along with the corresponding temperature and pain report. During each epoch, training and validation data were kept separate. For each potential mediator model, we performed a multi-level mediation analysis [69] on the test data and obtained p-values using a bootstrap approach with 5000 iterations. We chose the model with the most stable indirect effect for mediating thermal pain. Finally, we compared the results to those obtained using both mediation effect parametric mapping and the PDM approach.

For the HCP data, we combined our proposed algorithm with rCPM to find potential elements that mediate the relationship between fluid intelligence and accuracy of working memory task. We used 70% of the subjects for training and 30% for testing the model. The input data consisted of a set of vectorized connectivity values along with the corresponding fluid intelligence and working memory accuracy scores. We ran Steps 1-8 of the algorithm for 10 iterations. During each iteration, we used a 5-fold cross-validation grid search strategy on the training data to tune the model hyper-parameter. Thereafter, for each iteration, we fit the model with tuned parameters to the training data. Each iteration yields a potential mediator model, and similar to the analysis used for the pain data, we performed a multi-level mediation analysis on the test data to obtain p-values using a bootstrap procedure with 5000 iterations. Finally, we chose the model with the most stable indirect effect.

### E. Model interpretation

For both datasets, we used SHAP (SHapley Additive exPlanations) [70] to interpret the model fit. Shapley values are based on game theory which determines a ‘fair’ way to attribute the total gain to the players in a coalition game based on the individual contribution. The approach has recently been used to interpret deep learning models in a number of different medical applications [71–73].

In our application, the goal of SHAP is to explain the prediction obtained by the deep learning model by computing the relative contribution of each feature (e.g., voxel or connectivity edge) to the prediction. The Shapley values take into account the marginal distribution of every feature to the final prediction, making sure that the contributions of these features are optimally assessed. One drawback of using Shapley values is that they are computationally expensive. However, we used the Deep Shap implementation in python (https://github.com/slundberg/shap) which makes computation acceptable without compromising any inherent properties of the Shapley values.

## IV. RESULTS

### A. Simulations

Figure 3 shows the results of Simulation 1. Here we investigated how increasing the number of observations influenced the performance of our approach. We kept the dimensions of the mediator constant, but allowed the number of observations to vary. Clearly, as the number of observations increase the error bars become narrower, providing more accurate estimates of *α, β, γ*, and *αβ*. All three models perform roughly equivalently, though for small samples sizes the error bars for the deep learning model are somewhat larger, particularly when estimating *β*, indicating increased error variance. In contrast, the PDM approach shows a consistent bias in estimation of the *β* coefficient which leads to a slight underestimation (overestimation) of the indirect (direct) effect.

**FIG. 3.**
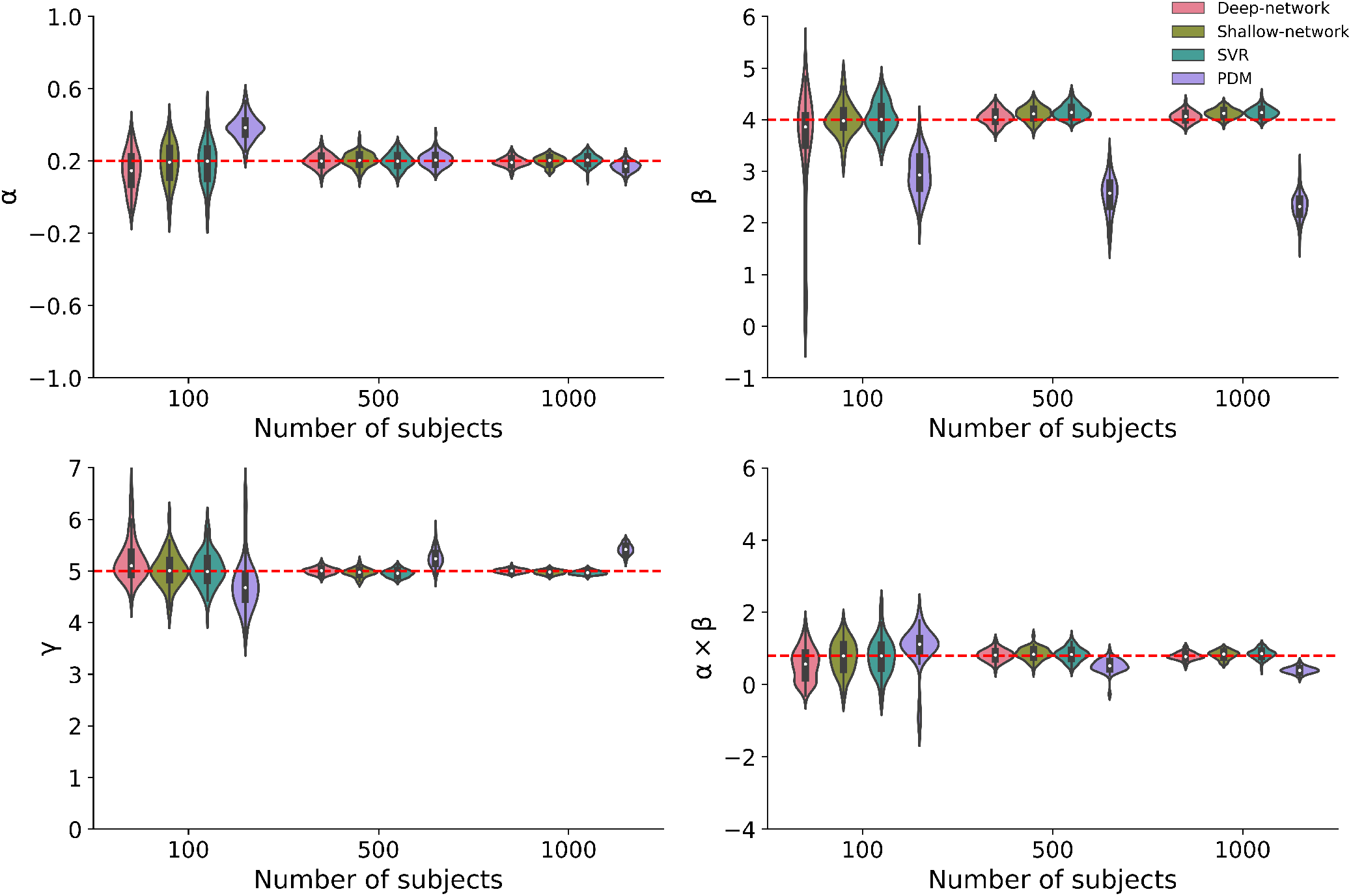
Results of Simulation 1, where we varied the number of observations (subjects) while keeping the dimensions of mediator constant. Each violin plot shows the estimated model parameters for the training dataset using deep learning model, shallow learning model and support vector regression. The red dotted line represents the ground truth data.

Figure 4 shows the results of Simulation 2. Here we investigated the ability of our approach to handle increased dimensions of the mediator variable. The values of all other variables remain the same as in Simulation 1. Again, all three models perform roughly equally. Interestingly, the error bars are constant across all dimensions. This indicates that the difficulty of the estimation problem is not directly related to the size of the mediator, but rather the information content which is constant as the images are simply scaled versions of one another. Again, the PDM approach shows a consistent bias in estimation of the *β* coefficient which leads to a slight underestimation (overestimation) of the indirect (direct) effect.

**FIG. 4.**
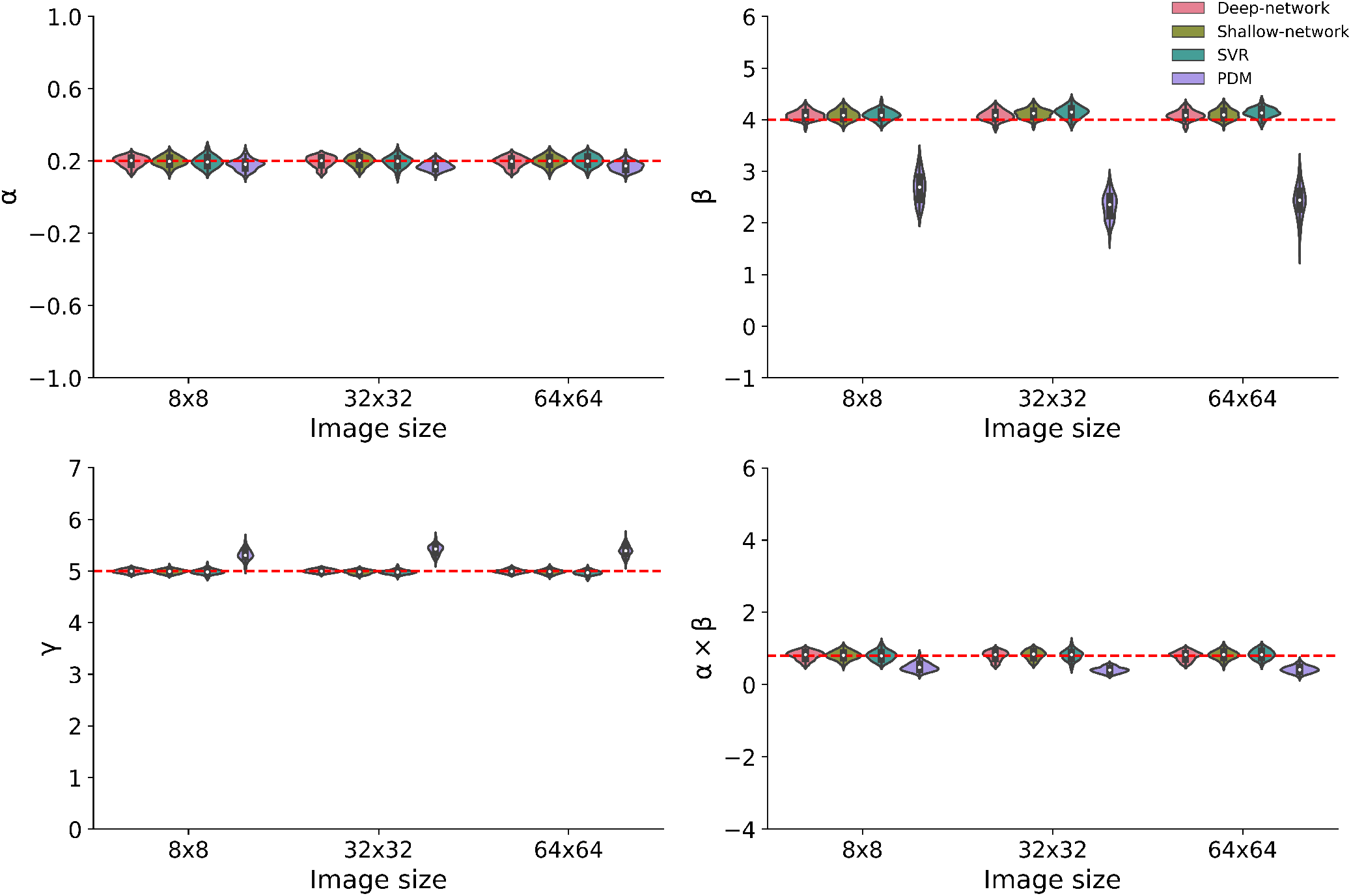
Results of Simulation 2, where we varied the dimension of mediator while keeping the number of observation constant (n=1000). Similar to simulation 1, each violin plot shows the estimated model parameters for the training dataset using deep learning model, shallow learning model and support vector regression. The red dotted line represents the ground truth data.

Figure 5 shows the results of Simulation 3. Here we investigated the performance of our approach in a ‘null’ setting where the indirect effect is 0. We used the same dimension of the mediator variables as described in Simulation 1, however we changed the value of *α* to be equal to 0. Each of the three models were able to handle this situation and on average found the indirect effect to be 0. Again, as the number of observations increase the error bars become narrower, providing more accurate estimates of *α, β, γ*, and *αβ*. Here, while the PDM approach shows a bias in estimation of both the *α* and *β* coefficients, both the direct and indirect effects appear unbiased.

**FIG. 5.**
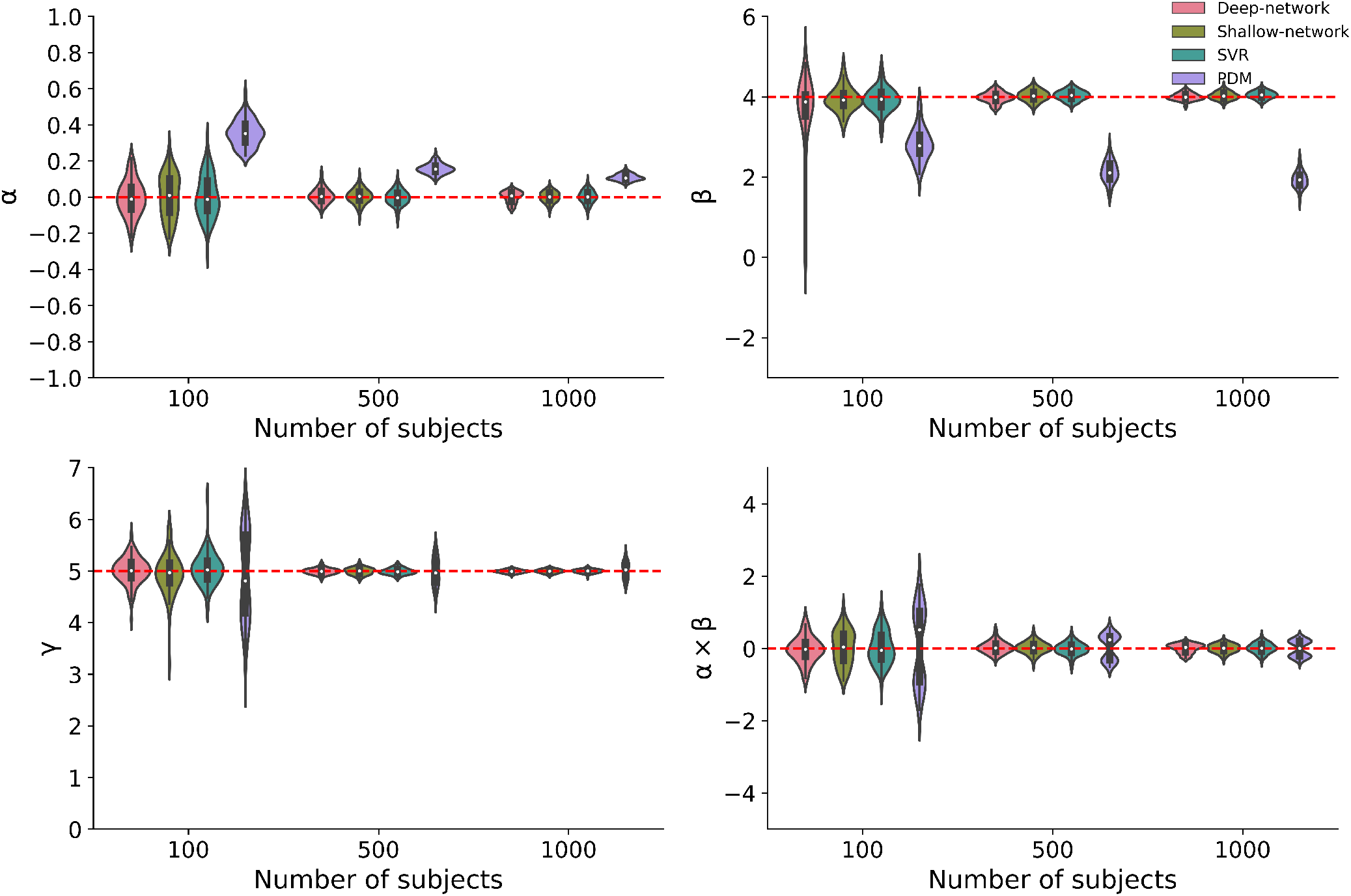
Results of Simulation 3, where we varied the number of observations (subjects) and removed the effect of mediation. Similar to simulation 1, each violin plot shows the estimated model parameters for the training dataset using the deep learning model, shallow learning model and support vector regression. The red dotted line represents the ground truth data.

### B. Pain Data

Algorithm 1 was combined with a deep learning algorithm and fit to the training data, consisting of 209 subjects with a total of 13372 trials from Studies 1-7. Each trial consisted of a temperature, a pain rating, and a 3-dimensional activation map. To validate the model fit, it was evaluated using an independent test dataset consisting of 75 subjects with a total of 2296 trials. Validation was performed by applying the trained deep learning model to the activation maps in the test dataset to obtain low-dimensional mediators. These were then placed into a standard three-variable path model together with the associated temperature and pain ratings. A multi-level mediation analysis [69] was performed on this data set, and the significance of *α, β*, and *αβ* was tested using a bootstrap procedure with 5000 iterations.

Figure 6A shows scatter plots illustrating the positive relationship between the low-dimensional mediator *z* and the input temperature, the pain ratings and the mediator, and the pain ratings and the temperature, respectively. Figure 6B shows the estimated *α* (stimulus intensity to brain path), *β* (brain to pain report path), and *αβ* (indirect) effects when applying the model fit to the training data. All results are significant (*p <* 0.05) when applied to the heat pain data, suggesting that the deep learning results are reliably related to pain and generalize across cohorts.

**FIG. 6.**
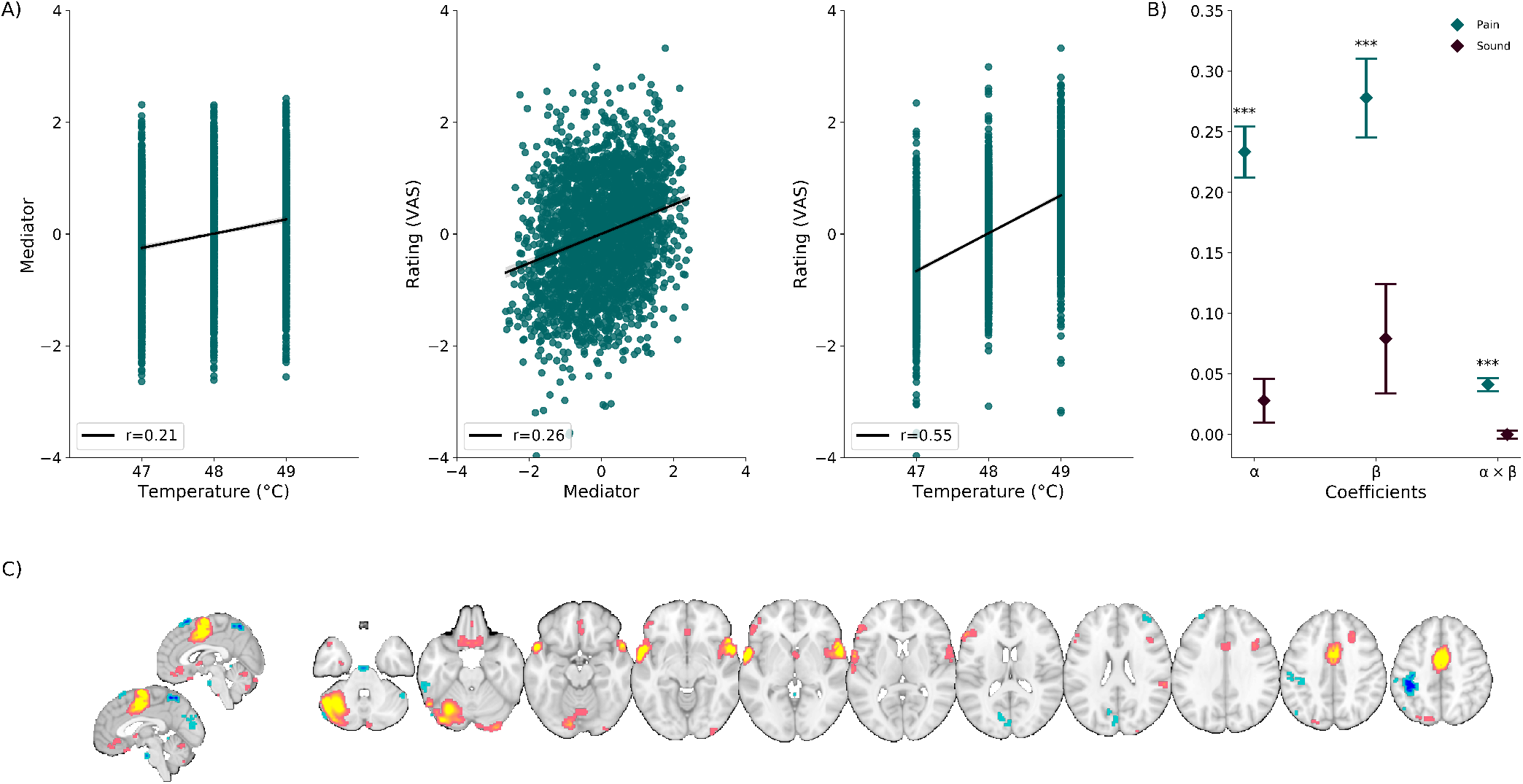
Validation on independent data (n=75). (A) Scatter plots show the relationship between the low-dimensional mediator and input temperature, pain ratings and mediator, and pain ratings and temperature of input stimuli, respectively. Lines show the least-squares fit between variables for each independent validation subject. (B) The estimated *α, β*, and *αβ* values obtained when applying the fitted machine learning model to the independent validation dataset, both for the pain and sound trials. These reflect the brain increases as a function of stimulus intensity, the relationship between brain and pain controlling for stimulus intensity, and the mediation effect, respectively. Error bars indicate SEM. *** *p <* 0.001. All coefficients were strongly significant for heat pain but non-significant for sound, indicating specificity to pain when compared with aversive sounds. (C) Voxel maps representing the 5% largest (in absolute value) Shapley values, indicating the regions involved in mediating the relationship between stimulus intensity and pain in the independent validation dataset. The majority of identified regions are targets of pain-related ascending pathways (e.g., somatosensory S1/S2, medial thalamus, Anterior Cingulate, and mid insular-opercular areas). Some regions are not generally considered to be related to primary pain pathways but play important modulatory roles (e.g., Ventromedial Prefrontal Cortex, Cerebellum, Anterior Temporal Cortices).

To determine which regions are driving the mediation, Shapley values were computed for all heat pain trials in the training data set. Figure 6C shows the voxels with the 5% largest absolute values. Brain regions shown are commonly associated with pain processing, such as multiple cerebellar regions, anterior cingulate and surrounding medial prefrontal cortex (MPFC), posterior medial orbito-frontal cortex (OFC)/ventromedial prefrontal cortex (vmPFC), lateral prefrontal cortex (area 47, inferior frontal sulcus [IFS], area 6), multiple temporal regions (temporal pole, TA2, entorhinal cortex), hippocampus, and Bed nucleus of Stria Terminalis (BST). It should be noted that the threshold was chosen arbitrarily, though it was determined that the maps were relatively stable on the range of 3-7%. Optimally, one could determine the significance of the regions that contribute to mediation effects using a bootstrap procedure. However, combining the Shapley analysis and bootstrap is computationally quite expensive in practice.

When considering the signs of the Shapley values, it is first worth noting that four different kinds of relationship are possible: (1) an increase in temperature leads to an increase in pain; (2) a decrease in temperature leads to a decrease in pain; (3) an increase in temperature leads to a decrease in pain; and (4) a decrease in temperature leads to an increase in pain. Here, type (1) is the standard, positive mediator case expected from nociceptive coding regions and type (2) represents a negative mediator, in which greater deactivation to stimulus mediates increased pain. Finally, types (3) and (4) are known as suppresor effects. Voxels shown in warm colors in Figure 6C correspond to those with positive values. As both *α* and *β* are positive, these regions represent positive mediators. They include brain regions commonly associated with pain processing, such as the dorsal posterior and mid-insula, S2, and MCC. Brain regions with negative weights represent negative mediators and are shown in cool colors, and include prefrontal regions, medial occipital, V1/V2/V3 and left sensorimotor cortex/parietal cortex, left S1/M1, parts of cerebellum, and the right amygdala/hippocampal border. Negative mediators are those that show less activation (or deactivation) with increasing temperatures, and lower regional activation is related to higher pain ratings. These types of relationships can be expected in brain regions whose function is inhibited by nociceptive input or that are deactivated with increased pain-related processing but are not considered as suppressor effects.

The test data also included trials with physically (e.g., knife on plate) and emotionally (e.g., screaming and crying) aversive sounds at three different pre-defined intensity levels. These trials were randomly intermixed with the heat pain trials. To test whether the results are specific for thermal pain, we applied the fitted model to these aversive non-painful stimuli. Application of the model fits on the sound data revealed no significant effects at the 0.05 significance level, see Figure 6B, indicating that the model does not mediate the relationship between sound intensity and intensity ratings. Thus, the results indicate a specificity to somatic pain compared to sound.

To further validate the findings we performed a number of follow-up analyses. First, we performed leave-one-study out cross-validation. Here we alternated which of the eight studies were used as the validation dataset, while training on the remaining seven studies. Results can be seen in Figure S3. In total five of the eight pain datasets were significant when used as the test dataset. Interestingly, the three studies that were not significant (EXP, IE, and SCEBL) are the ones with the strongest psychological interventions, and the effect of pain depends strongly on these interventions. For EXP and IE, there are cues prior to pain stimulus that state whether high or low pain is coming. For SCEBL there is a cue that states how other subjects responded to the upcoming stimuli. Much of the pain response is likely linked to these cues, and therefore in each case it is not entirely surprising that the *β*-pathway is non-significant. Second, we also ran multiple iterations of *k*-fold cross validation. Due to computational constraints we restricted the number of replications to 3 times. During the *k*-fold cross validation, the training dataset is split into 3 folds. Figure S4 shows the estimated *α, β* and *αβ* values obtained when applying the fitted machine learning model to the left-out fold for the pain trials. As seen in the figure, all coefficients were strongly significant.

Next, we compared the results with those obtained using two competing approaches: the PDM approach and a mass univariate mediation effect parametric mapping approach. In Figure S5 we show significant voxels obtained through both analyses. Both maps are thresholded at a false discovery rate (FDR) of *q <* 0.05. The PDM approach linearly combines information across images into a smaller number of orthogonal components that are chosen based on the proportion of the indirect effect that they explain. Here we only use the first PDM which corresponds to the linear combination that maximizes the proportion of the indirect effect explained. Similar to the proposed approach, the PDM approach found mid insular-opercular areas, somatosensory S1, S2 and medial thalamus mediated the temperature-pain relationship; see Figure S5(a). In contrast, mediation effect parametric mapping fits an independent mediation model on each individual voxel in the fMRI data. Thereafter, brain regions corresponding to the intersection of voxels with significant paths *α, β* and *αβ* are interpreted as mediators. The mass univariate analysis found the cerebellum, posterior and midinsula, MCC, S2 and S1 were significant mediators; see Figure S5(b). Comparing these results to the proposed approach found both similarities and differences. For example, both maps included somatosensory regions of MCC, mid insula, S2 and cerebellum. Additionally, negative mediators in prefrontal regions, medial occipital, S1 and right amygdala/hippocampal border region were not identified by the univariate mediation model.

### C. HCP data

Algorithm 1 was combined with a ridge regression connectome-based predictive model and fit to the training data, consisting of 558 subjects. Each subject’s data consisted of fluid intelligence, a 1-dimensional vectorized functional connectivity matrix, and a working memory accuracy score. To validate the results, they were applied to a test dataset consisting of 240 subjects. Validation was performed by applying the trained rCPM model to the elements of the functional connectivity matrices in the test data to obtain low-dimensional mediators. These were then placed into a standard three-variable path model together with the associated fluid intelligence and accuracy scores. A multi-level mediation analysis [69] was performed on this data set, and the significance of *α, β*, and *αβ* was tested using a bootstrap procedure with 5000 iterations. Note that in practice, the mediation analysis could have been run in either direction (i.e., with working memory accuracy as the *X* variable and fluid intelligence as the *Y* variable). Hence, there is no strong causal interpretation to be made here, but rather this is an example of mediation analysis can identify brain patterns jointly related to two variables that are part of the same system.

Figure 7 shows the results of applying the fitted rCPM mediation model to a test data set from the HCP dataset. Figure 7A shows scatter plots illustrating the positive relationship between the low-dimensional mediator and fluid intelligence, accuracy and the mediator, and accuracy and fluid intelligence, respectively. Figure 7B shows that the effects are significant in the test dataset, indicating that the functional connectivity matrix mediates the relationship between fluid intelligence and working memory performance (accuracy) in a manner that generalizes across cohorts.

**FIG. 7.**
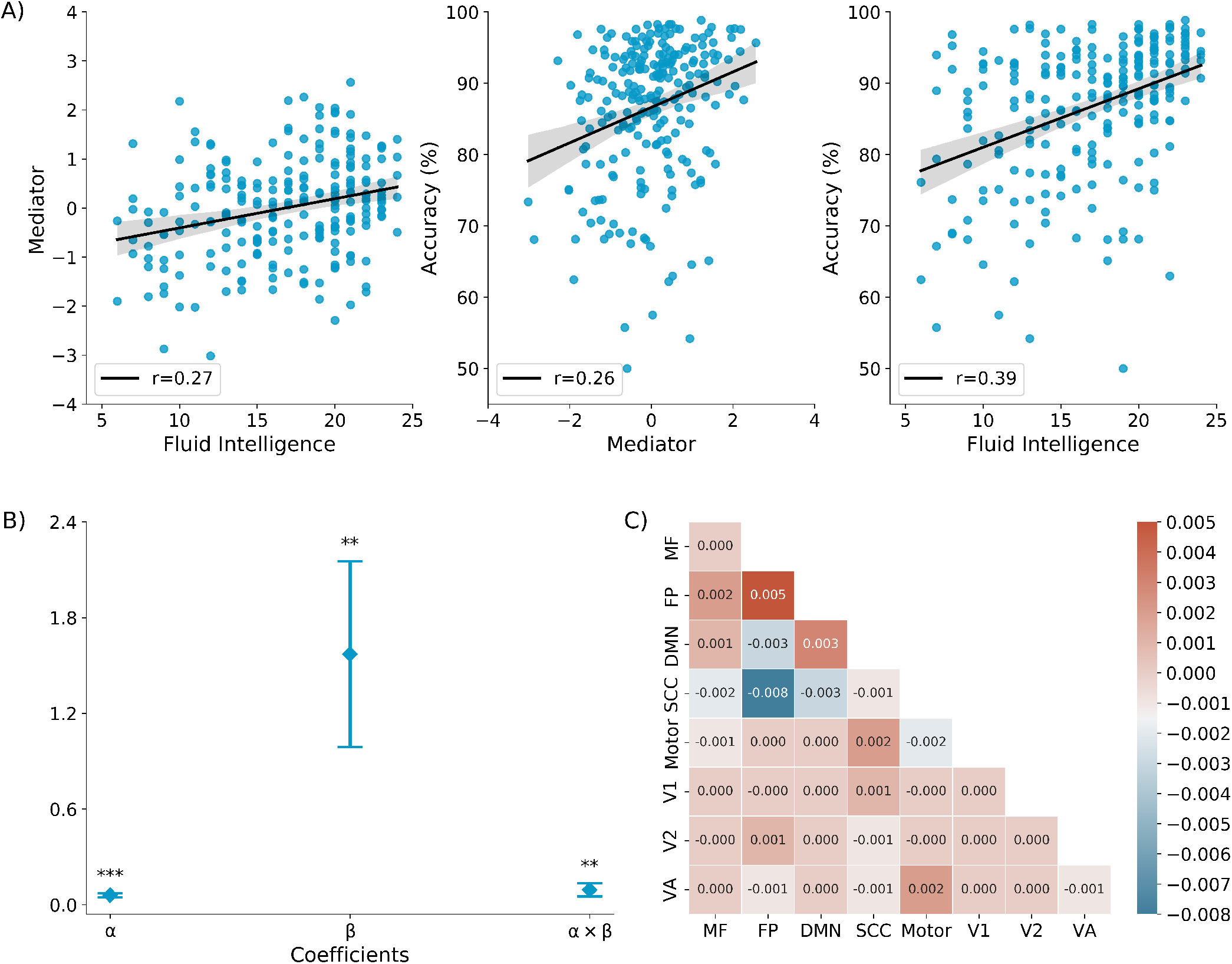
Results on test data (n=240). (A) The high dimensional functional connectivity matrix which serves as a mediator (*M*) of the relationship between fluid intelligence (*X*) and the accuracy on the working memory task (*Y*). (B) Scatter plots show the relationship between the low-dimensional mediator *z* and fluid intelligence, accuracy and the mediator, and accuracy and fluid intelligence, respectively. Lines show the least-squares fit between variables for each test subject (*n* = 240). (C) Shapley values averaged over voxels connecting each pair of large-scale networks. High connectivity within the Frontoparietal (FPN) and Default Mode Networks (DMN), and low/negative connectivity between FPN and DMN and between FPN and subcortical regions (among other connections) mediated the relationship between fluid intelligence and accuracy of working memory task. This demonstrates links between brain connectivity and both working memory performance and fluid intelligence. MF: Medial frontal, FP: Frontoparietal, DMN: Default model network, SCC: Subcortical-cerebellum, V1: Visual I, V2: Visual II, VA: Visual association. *** *p <* 0.001 and ** *p <* 0.01.

Next, we determined the SHAP values in order to determine which connections are driving the mediation. Figure 7C shows the SHAP values in the test dataset averaged over subjects and components in each of seven pre-defined networks [74]. Connections with positive weights are shown in warm colors. As both *α* and *β* are positive, these connections represent positive mediators. They include brain networks such as the Frontoparietal and Default Model Network, and connectivity between Frontoparietal and Medial Frontal Networks, Motor and Subcortical-Cerebellum Networks, and Motor and Visual Association Networks. Connections with negative weights represent negative mediators and are shown in cool colors. They include the motor network, and connectivity between Frontoparietal and Subcortical-Cerebellum Networks, Frontoparietal and Default Mode Networks, and Default Mode and Subcortical-Cerebellum Networks. The negative weights indicate that these connections show lower values with increasing fluid intelligence, and that lower connectivity is related to higher working memory accuracy.

## DISCUSSION

In this work we introduce a novel analytic approach for identifying high dimensional mediators that links exposure variables, high-dimensional brain measures, and behavioral outcomes into a single unified model. Using the approach, the effects of the exposure on the outcome are decomposed into separable direct and indirect effects, representing the influence of the variables *x* on *y* unmediated and mediated by *m*, respectively. The indirect effect is determined by the product of the coefficients *α* and *β*, while the direct effect is determined by the coefficient *γ*; see Figure 1 for more detail. Our approach is flexible, allowing for easy plug-and-play with different machine learning models depending on the type of data being analyzed. We demonstrate this flexibility in two applications, that necessitate using two different classes of machine learning models.

In the pain application, we used the proposed approach together with a deep learning model to identify brain networks that mediate the relationship between stimulus intensity and pain reports. To interpret the results and determine which regions mediated the temperature-rating relationship we computed maps of Shaply values; see Figure 6. We arbitrarily chose the largest 5% Shapley values when presenting our findings, but found that the maps were relatively stable across a range from 3-7%. Optimally, one would determine the significance of the regions that contribute to mediation effects using a bootstrap procedure. However, combining Shapley analysis and the bootstrap is extremely computationally expensive in practice.

Importantly, the derived mediators generalized to independent pain data, but not to aversive sound data, which indicates a degree of specificity of the model for pain. Several previous studies [29, 58] have identified brain mediators of pain in a univariate manner by investigating each voxel separately. A shortcoming of this approach is that it can potentially miss brain regions whose contributions to pain perception are conditional on other regions. In addition, researchers have found that functional information in the brain is likely encoded in distributed patterns across neural ensembles and systems [75, 76]. This implies that brain information should ideally be treated in a multivariate fashion [77, 78], highlighting the importance of using multivariate brain mediators to characterize these patterns. Thus, we believe our approach provides a more comprehensive picture of pain processing in the human brain than studies that use univariate analyses, or focus solely on the stimulation-brain or brain-outcome relationships.

It should be noted that the pain data was previously analyzed using the principal directions of mediation (PDM) approach [35], which is an alternative method for performing high-dimensional mediation developed by our group. As both the machine learning-based approach and the PDM approach seek to estimate distributed, network-level patterns that formally mediate the relationship between stimulus intensity and pain, this allows for a convenient comparison between methods. The PDM approach linearly combines activity in different mediators into a smaller number of orthogonal components, with components ranked based upon the proportion of the indirect effect that each accounts for. In contrast, the proposed approach provides a non-linear combination of mediators as defined by the deep learning architecture. The results obtained using both methods are roughly equivalent, with both methods highlighting the same regions as mediators and providing results specific for pain vs. aversive sounds. Using the PDM results as a benchmark, we believe this provides evidence of the efficacy of our new machine-learning based approach. That said, the proposed approach has several benefits over the PDM approach. One is the aforementioned ability to study non-linear combinations of the original high-dimensional mediators. Another is its flexibility to be applied to a wide array of different data, for example, brain connectivity data.

In this application we used a deep learning model to investigate its ability uncover brain regions that mediate the relationship between temperature and pain rating. In general, we believe that a simpler machine learning approach is preferable when the more complicated models do not show empirical evidence for improvement. Therefore, for completeness we repeated the analysis using both SVR and Ridge regression in place of the deep learning model. We found that the deep learning model outperformed both SVR and Ridge regression. Moreover, we did not obtain interpretable results when studying the model weights for either SVR and Ridge regression, even though both are linear models.

In the application to HCP data, we used the proposed algorithm together with a connectome-based predictive model [37] to find elements of the resting-state connectivity matrix that mediate the relationship between fluid intelligence and working memory accuracy. The link between fluid intelligence and working memory capacity has long been established [79, 80]. In recent work, [81] fit separate connectome-based predictive models to predict working memory performance and fluid intelligence, respectively, from whole-brain functional connectivity patterns observed in HCP participants. They found that overlap between the working memory and fluid intelligence networks were limited to connections between prefrontal, parietal, and motor regions. Additionally, [82] have found that activity in “multiple demand” networks (i.e. lateral and dorsomedial frontal areas, anterior insular areas, and areas along the intra-parietal sulcus regions) was robustly associated with more accurate and faster responses on a spatial working memory task and fluid intelligence. Our approach extends this approach by providing a unified model that links working memory accuracy, fluid intelligence, and functional connectivity. Using our approach, we found the strongest connections within Frontoparietal, Default Mode, and Motor networks, and between Frontoparietal, Default Mode, and Subcortical-Cerebellum networks. This evidence aligns well with findings from lesion studies that have also reported a selective relationship between fronto-parietal regions and working memory task as well as fluid cognitive abilities [83, 84]. However, it is a further challenge to identify and interpret if these connections are statistically significant in mediating the relationship between fluid intelligence on working memory accuracy.

Interpreting the indirect effect is an important part of mediation analysis. The proposed high-dimensional mediation approach can be placed into a potential outcome framework to access the conditions necessary for causal mediation analysis. In short, using potential outcomes notation, let *M* (*x*) denote the value of the mediators if treatment *X* is set to *x*. In our example, this represents the brain activation corresponding to a temperature set to a particular value *x*. Similarly, let *Y* (*x, m*) denote the outcome if *X* is set to *x* and *M* is set to *m*. This is the reported pain corresponding to both temperature and brain activation set to *x* and *m*, respectively. Using this notation, the natural unit indirect effect can be defined as *Y* (*x, M* (*x*)) −*Y* (*x, M* (*x*^*^)). This corresponds to the change in pain rating that arises when brain activation is switched from *M* (*x*) to *M* (*x*^*^). The *αβ*-effect represents the average indirect effect, which is equivalent to the natural direct effect when there is no treatment-mediator interaction. In other words, when *M* (*x*) and *Y* (*x, m*) are well defined and a series of assumptions hold, *αβ* can be used to identify causal mediation effects. In practice, it is difficult to test whether these assumptions hold. Hence, we refrain from any causal interpretations of our results in this work. This material is discussed in the context of high-dimensional mediation in greater detail in earlier work by our group [30, 34].

Though our proposed framework is versatile and provides an option to test any number of machine learning models to find mediators using high dimensional data, it has its limitations. For example, the outcome of our framework depends on the performance of the underlying machine learning model. This implies that one needs to build a model that is able to accurately represent the relationship between the high dimensional mediator and the outcome. A failure to yield an expected result might be linked to a poor model selection and one needs to be careful before drawing conclusions especially in clinical applications. It should be noted that problems associated with building a good machine learning model for predicting outcome is an overall challenge for the entire field that is not unique to the proposed method. In addition, there is reason to believe that there are situations where prior knowledge about the data or its acquisition plays an important role in the mediation analysis. For instance, prior knowledge about the brain function and structure couldbe a crucial factor in constraining mediation analysis. In our initial implementation, we have not considered such prior knowledge, but these factors can be incorporated into the machine learning model and thus utilized in our approach. We leave this for future research.

In conclusion, we have developed a new approach for identifying high dimensional mediators. Our proposed method provides a potential way for overcoming challenges with finding mediators in high dimensional data. Our single unified deep learning method reduces the high dimensional mediator to a single latent intermediate mediation measure. Such a measure can be used to study how dimensional mediators mediate the relationship between various traits and be applied to a variety of clinical applications. We applied our method to two different types of data, thus illustrating the robustness of the method. The development of methods for dealing with high dimensional mediation is in its infancy and this is the first application of deep learning to the field.

## ACKNOWLEDGEMENT

Data were provided by the Human Connectome Project, WU-Minn Consortium (Principal Investigators: David Van Essen and Kamil Ugurbil; 1U54MH091657), which was funded by the McDonnell Center for Systems Neuroscience at Washington University and the 16 NIH Institutes and Centers that support the NIH Blueprint for Neuroscience Research. This research was supported in part by NIH grants R01EB016061 and R01EB026549 from the National Institute of Biomedical Imaging and Bioengineering, R01MH076136 from National Institute of Mental Health, and Oracle Cloud credits and related resources provided by the Oracle for Research program. We are particularly thankful to Bryan Barker and Rajib Ghosh for providing support with the Oracle cluster.

## V. SUPPLEMENTAL MATERIAL

**FIG. S1.**
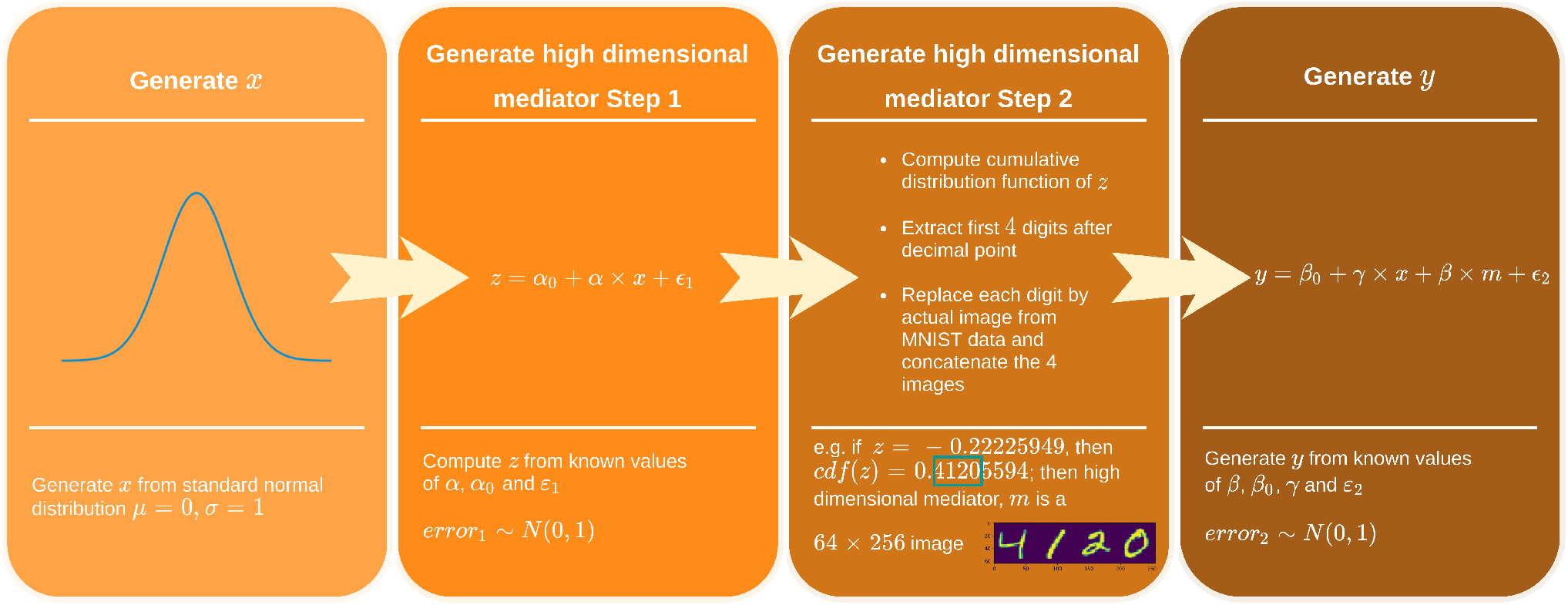
Overview of simulation. Generate *x* from a standard normal distribution. For known values of *α*_0_, *α* and *ϵ*_1_, compute *z* using *z* = *α*_0_ + *αx* + *ϵ*_1_. In order to generate high dimensional mediators *m*, first, compute the *cumulative distribution function of z* (i.e., *norm*.*cdf* (*z*)). Thereafter, take the first 4 digits after the decimal point and replace these digits by images from the MNIST data [47]. Concatenate the four images to create a high dimensional mediator. For known values *β*_0_, *β, γ* and *ϵ*_2_ generate *y* using *y* = *β*_0_ + *γx* + *βz* + *ϵ*_2_. The image size of MNIST data can be re-scaled to modify the overall dimension of the mediator.

**FIG. S2.**
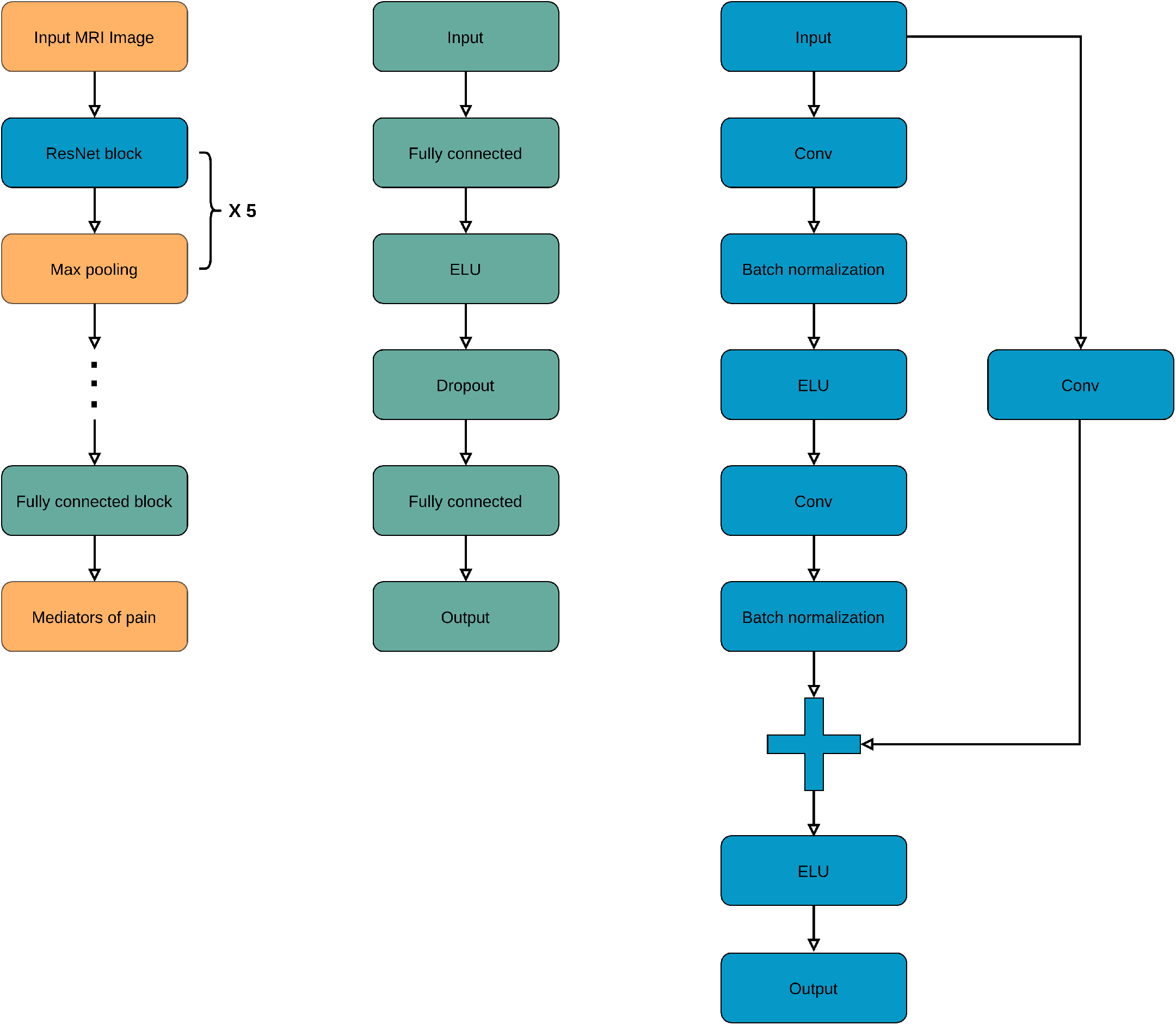
A flowchart of the deep learning model architecture. The input to the model is the single trial brain activation maps. Thereafter, the architecture comprises a series of ResNet and max pooling blocks. Finally, it has a fully connected block which eventually predicts the mediators of pain. The backbone of the deep learning model architecture is the residual block which is concatenated to a max pooling layer. Together these two blocks are repeated five times and the output of the last block is concatenated to a fully connected block. The fully connected block consists of a fully connected layer with an exponential linear unit (ELU) activation function. Its output is connected to a dropout layer [85] which randomly drop units from the neural network during training in order to avoid over-fitting. The ResNet block consists of a 3-D convolutional layer with kernel size 3×3×3 and stride 1×1×1 followed by a batch normalization layer and an ELU activation function. The convolutional layer and batch normalization layers are repeated twice and concatenated to the output of skip connection.

**FIG. S3.**
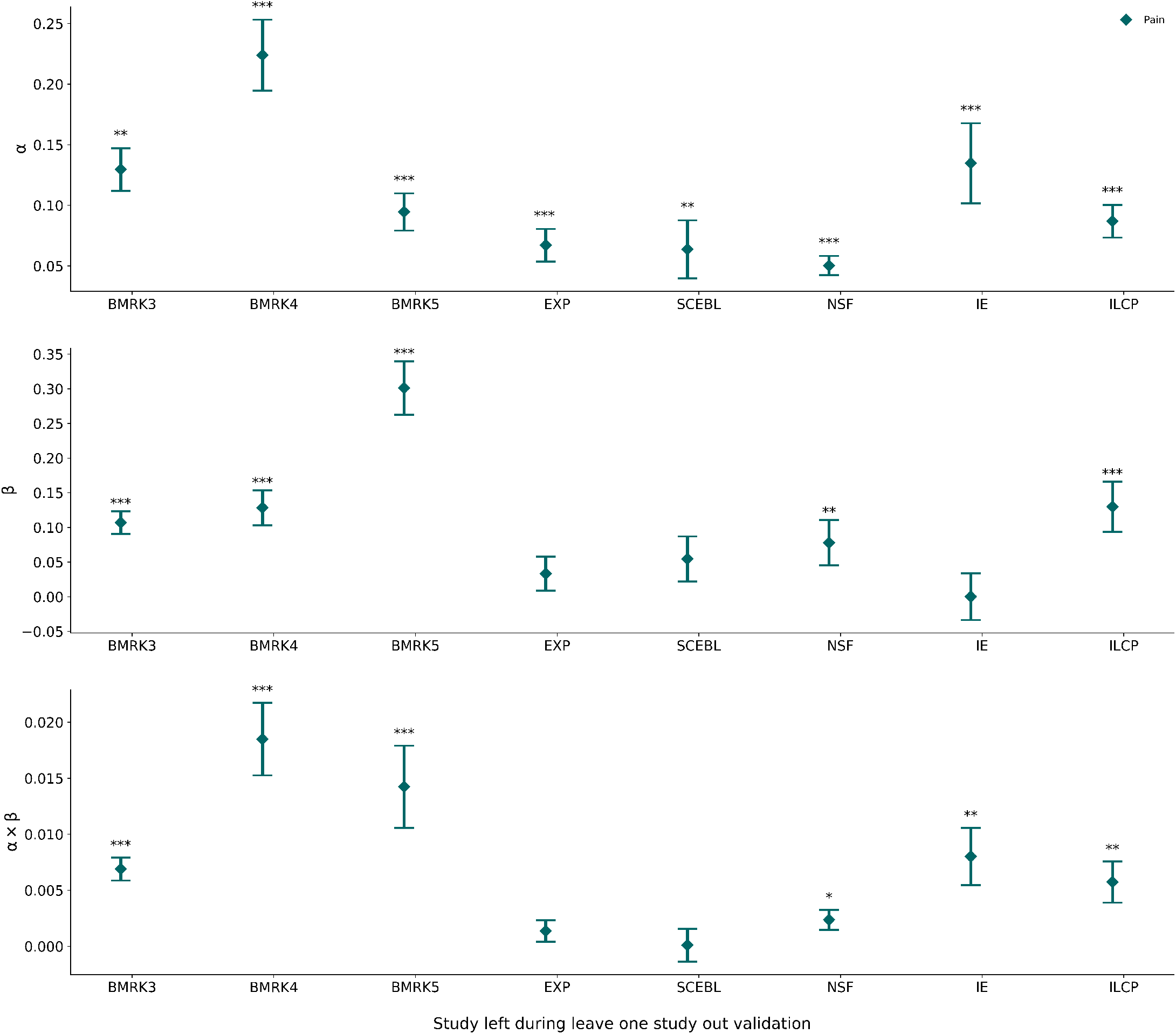
Result on a leave-one-study-out validation. During the leave-one-study-out validation, each study was left out during training the model and used only during testing the model. The estimated *α, β* and *αβ* values obtained when applying the fitted machine learning model to the left out study dataset for the pain trials. Error bars indicate SEM. *** p < 0.001. In total five of the eight pain datasets are significant when used as the test dataset. The three studies that were not significant (EXP, IE, and SCEBL) are the ones with the strongest psychological interventions, and the effect of pain depends strongly on these interventions. For EXP and IE, there are cues prior to pain stimulus that state whether high or low pain is coming. For SCEBL there is a cue that states how other subjects responded to the upcoming stimuli. Much of the pain response is likely linked to these cues, and therefore in each case it is not entirely surprising that the *β*-pathway is non-significant.

**FIG. S4.**
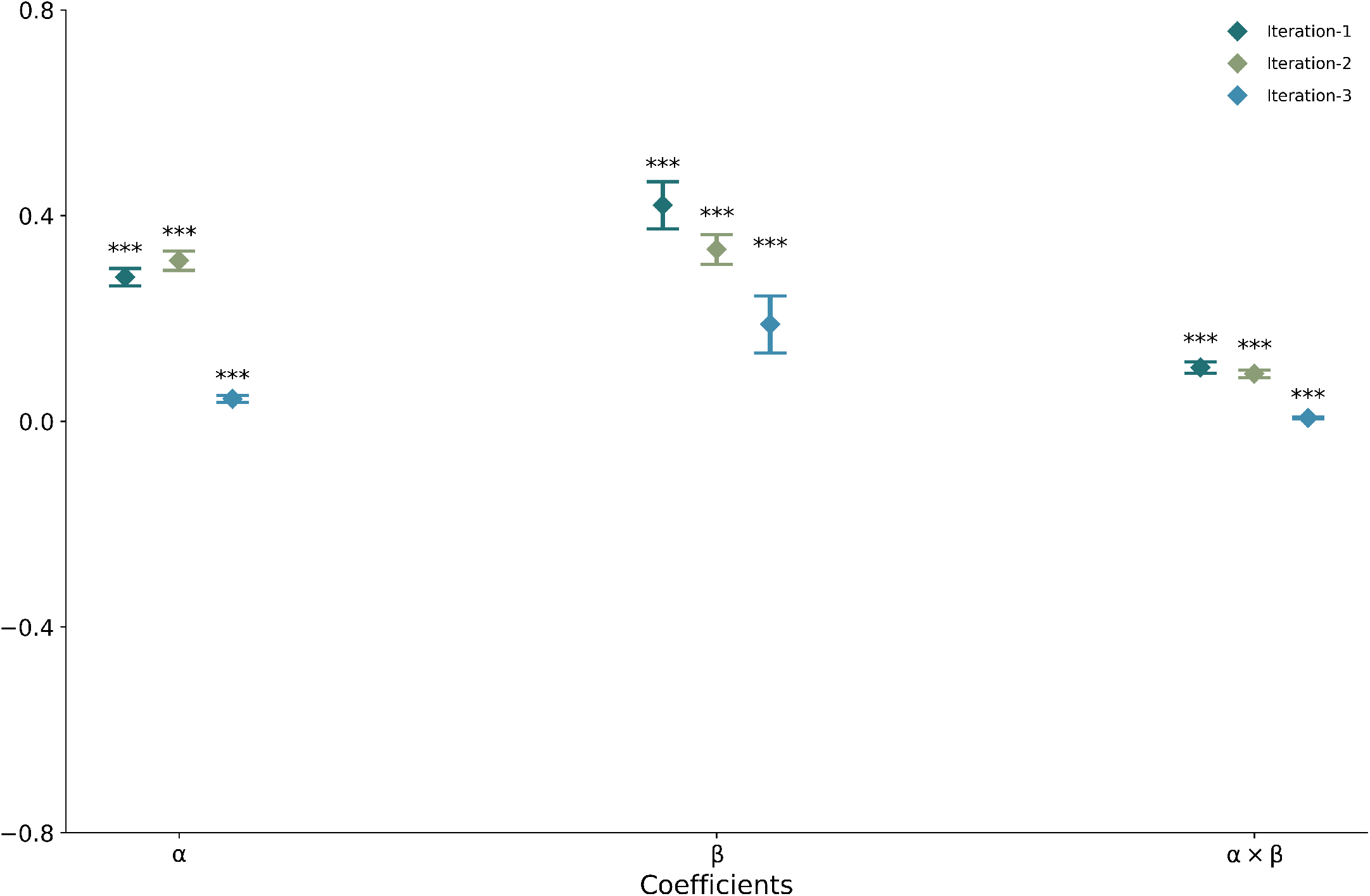
Results from multiple iterations of *k*-fold cross validation. Due to computational constraints we restricted the number of replications to 3 times. During the *k*-fold cross validation, the training dataset is split into 3 folds. The estimated *α, β* and *αβ* values obtained when applying the fitted machine learning model to the left out fold for the pain trials. Error bars indicate SEM. *** p < 0.001. All coefficients were strongly significant for pain trials.

**FIG. S5.**
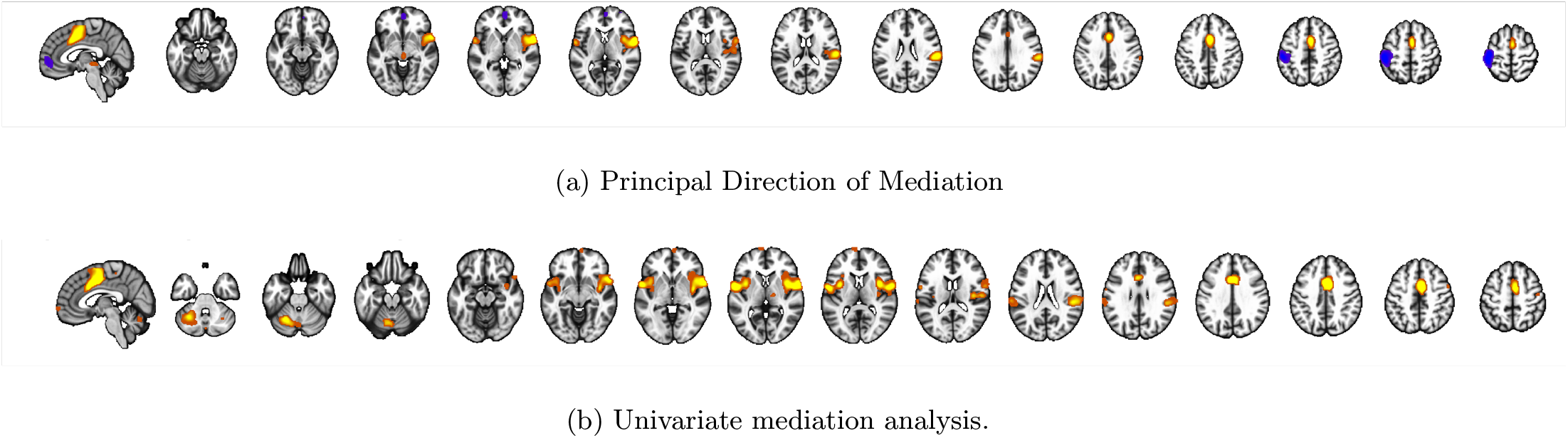
Comparison with PDM and a mass univariate approach (mediation effect parametric mapping) approach. Panels (a) and (b) show maps with individually significant voxels at FDR *q <* 0.05 from a PDM and univariate mediation analysis, respectively. These results were previously discussed in [35].

